# Viral Infection Skews Immunoglobulin Light Chain Repertoire Diversity

**DOI:** 10.1101/398529

**Authors:** John C. Schwartz, Michael P. Murtaugh

## Abstract

Antibody responses are fundamentally important to effector and memory mechanisms of disease resistance. Antibody repertoire diversity and its response to natural infection is poorly understood, yet is a prerequisite for molecular and structural elucidation of functionally protective immunity to viral infections. Using a swine model of mammalian viral infection, we observed marked changes following infection with the major porcine pathogen, porcine reproductive and respiratory syndrome virus (PRRSV). Deep sequencing of >516,000 light chain VJ mRNA genes showed that, similar to humans, swine utilize both lambda and kappa loci equivalently. However, V and J gene usage were highly restricted; ≥99% of lambda light chains were *IGLV3* and *IGLV8* family members joined to *IGLJ2* or *IGLJ3*, and 100% of kappa locus transcripts were *IGKV1* or *IGKV2* with only *IGKJ2*. Complementarity-determining region (CDR) variation was limited. Nevertheless, total diversity richness estimates were 2.3 × 10^5^ for lambda and 1.5 × 10^5^ for kappa, due in part to extensive germline variation in framework regions and allelic variation. Infection by PRRSV reduced total richness due to expression of several highly abundant clonal populations. Antibody light chain repertoires differed substantially among individuals, thus illustrating extensive potential variation in immune response in outbred populations. These findings demonstrate that individual variation in light chain repertoires may be an important component of variable antibody responses to infection and vaccination, and that swine are a relevant model of human antibody diversification in which the immune response capacity is critical to understanding individual variation in immune protection against disease.

**Conflict of interest statement:** The authors declare no conflicts of interest.

**Highlights:**

- λ and κ light chain diversity is equivalent to heavy chain diversity
- High diversity is present despite limited gene segment usage
- PRRSV infection increases abundance of dominant λ and κ VJ clones
- High levels of variation are present among animals

## Introduction

Outbred populations of humans and animals show extensive individual variation in the efficacy of immune responses to natural infections and vaccinations, even though genetic and biochemical mechanisms used to generate diversity within individual animals have an enormous capacity for recognition of antigenic structures. For example, the BCG (Bacillus Calmette–Guérin) vaccine provides a wide range of protection against tuberculosis in humans, ranging from nearly ineffective to nearly completely effective (1, 2). Vaccination of pigs against foot and mouth disease virus (FMDV) is characterized by significant animal-to-animal variation in anti-FMDV IgA (3); and post-vaccination anti-FMDV antibody responses in cattle are significantly affected by genetic background (4). Furthermore in swine there is substantial antibody response variation to vaccination against *Actinobacillus pleuropneumoniae* and to infection with *Mycoplasma hyorhinis* (5, 6). While many factors may account for individual variation in the immune response to an antigen, the size of the antibody repertoire and variation in repertoire composition could be significant host-dependent factors.

Swine are susceptible to a range of bacterial and viral contagions similar or identical to those in humans. As in other species, the porcine immunoglobulin genes are organized in three loci: a heavy chain locus on chromosome 7 and kappa (IGK) and lambda (IGL) light chain loci on chromosomes 3 and 14, respectively (7). We previously characterized the genomic organization of the kappa and lambda light chain immunoglobulin loci (8, 9). Within the kappa locus, there are three functional *IGKJ* genes and the first 14 *IGKV* genes have been identified, of which nine are putatively functional (8). The porcine lambda locus possess two functional *IGLJ* genes and 12 putatively functional *IGLV* genes belonging to three families, of which the *IGLV3* and *IGLV8* families comprise the vast majority of known expression (9, 10). We also previously identified substantial allelic variation among the germline *IGLV* (11) and described *IGLV3-6* which is present in some animals as either a pseudogene or as a functional highly expressed gene (12).

To further analyze the diversity of the light chain repertoire and its response to disease challenges, we sequenced and analyzed antibody light chain amplicon libraries obtained from healthy and virally-infected pigs. The analyses confirmed limited combinatorial diversity, yet extensive complementarity-determining region (CDR) and framework variation, likely arising from somatic hypermutation. Marked changes in abundance and repertoire diversity were observed following porcine reproductive and respiratory syndrome virus (PRRSV) infection that were indicative of clonally amplified and potentially PRRSV-specific B cells. Our observations also suggest that the utilization (or lack thereof) of *IGLV3-6* may skew the antibody repertoire toward expressing other genes at higher levels.

## Materials and Methods

### Samples

The animal studies were reviewed and approved by the University of Minnesota Institutional Animal Care and Use Committee. Lymphoid tissues (spleen, palatine tonsil, inguinal lymph node, and bronchial lymph node) from five genetically-similar, commercially-sourced piglets were obtained from a previous study (13). Piglets were challenged with virulent PRRSV strain JA142 (n=3) or not (n=2) at 3-4 weeks of age and tissues were harvested 63 days after infection and stored in RNAlater (Ambion/Thermo-Fisher). Tissues were homogenized and total RNA was extracted using the RNeasy Mini Kit (Qiagen) and reverse transcribed using the QuantiTect Reverse Transcription Kit (Qiagen).

### Library generation and sequencing

Primers were designed in Primer3 set to default parameters (14). *IGKV* and *IGLV* gene family-specific forward primers were designed using the conserved leader region sequences from annotations of the porcine kappa and lambda loci. In order to detect possible functional alleles of previously annotated pseudogenes, forward primers were designed such that all known kappa and lambda V genes could be amplified from cDNA regardless of functionality. Hence, the leader region for *IGLV8-21*, a pseudogene, was distinct enough to warrant a separate forward primer (Table 1). Reverse primers were designed to be specific for *IGKC* and the three nearly identical *IGLC* genes (8, 9).

**Table 1.**
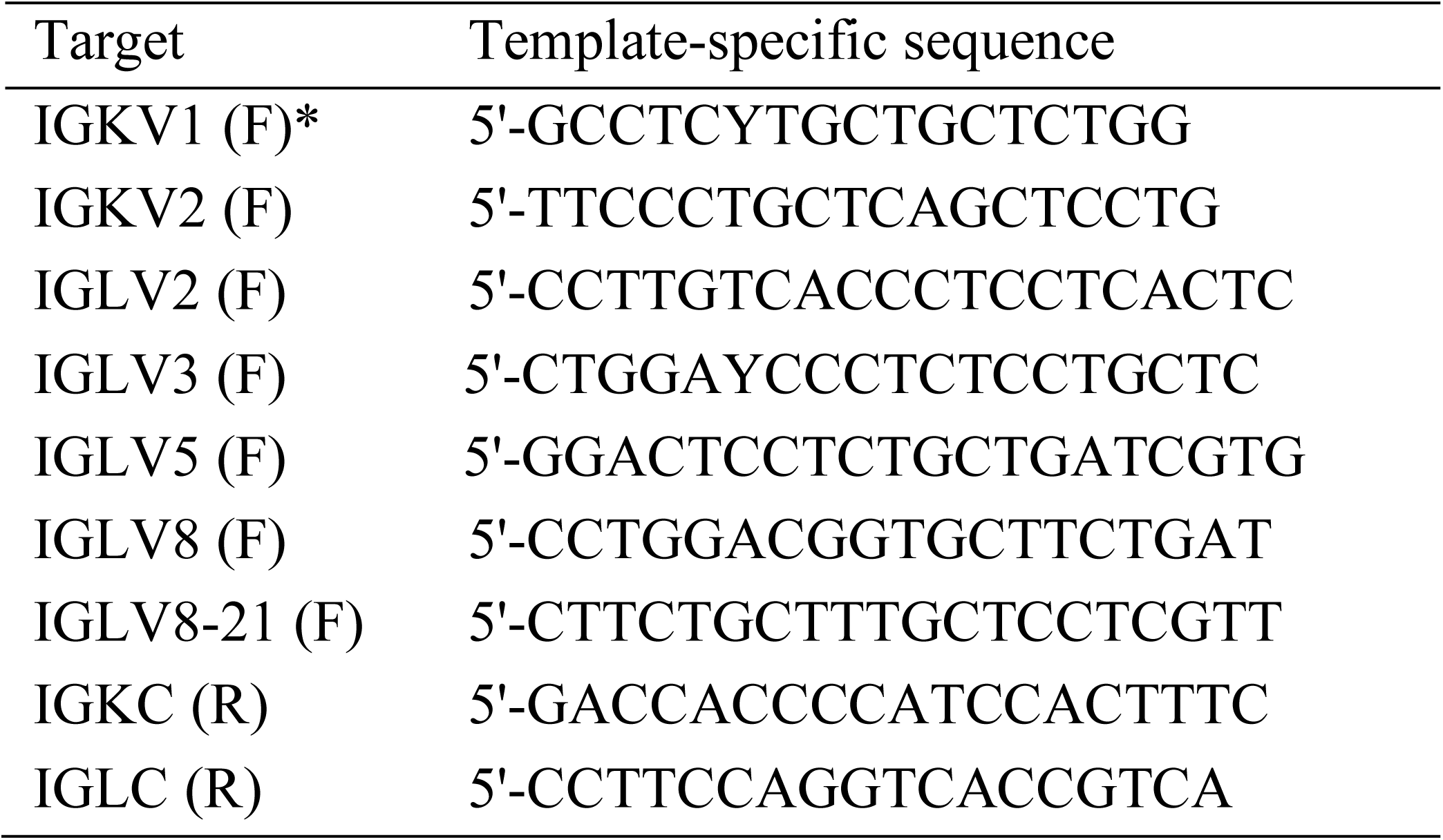
Primers used for amplification of porcine light chain cDNA VJ rearrangements.

The Roche 454 Titanium fusion adapter A (5′-CGTATCGCCTCCCTCGCGCCATCAG) was added to the 5′ end of each forward primer and fusion adapter B (5′-CTATGCGCCTTGCCAGCCCGCTCAG) was added to the 5′ end of each reverse primer. Ten base pair molecular ID tags were added to the forward primer between the fusion adapter and the template-specific sequence to identify animal source. PCR reactions were performed separately by tissue and products were pooled according to pig, quantified, and re-pooled in equimolar amounts so that the final library contained approximately the same number of amplicons from each animal. Pooled amplicons were gel-purified using the QIAquick gel extraction kit (Qiagen). To remove trace amounts of agarose, the amplicons were purified again using the QIAquick PCR purification kit (Qiagen). Roche Titanium 454 pyrosequencing was performed at the W. M. Keck Center for Comparative and Functional Genomics, University of Illinois at Urbana-Champaign.

### Sequence analysis

Pyrosequencing reads were deposited in the National Center for Biological Information (NCBI) sequence read archive (SRA) database under accession number SRP026119 (BioProject: PRJNA206406). Short reads (< 350 bp), reads with a missing or aberrant molecular ID tag, and reads containing premature stop codons were excluded from further analyses. *IGKV* and *IGLV* gene usage was determined using BLAST (15). Reads were translated into putative amino acid sequences using EMBOSS within the Galaxy framework (16, 17). For diversity analysis, amino acid sequences were first clustered using CD-HIT at an identity threshold of 100% (18) and these clusters were used to generate abundance curves and richness estimates. CD-HIT and Chao1 were used to calculate lower-bound estimates of repertoire richness (19-21). The clustered results were used to generate abundance curves within Microsoft Excel. Total number of sequence clusters and the numbers of singletons and doubletons generated from CD-HIT output were used to calculate Chao1 lower-bound estimates of repertoire richness (19-21).

## Results

### Sequencing results

A total of 516,097 pyrosequencing reads were obtained with a mean length of approximately 510 bp and a median length of 521 bp. Reads shorter than 350 bp which did not span the entire variable region were removed from further analyses (79,172 reads). An additional 11,739 reads with aberrant molecular ID tags and 53,046 reads containing premature stop codons and/or ambiguous amino acids were also removed from further analyses. Thus, a total of 372,140 full-length, in-frame reads were analyzed for V and J gene usage. The percentages of reads from individual pigs were approximately equal (pig 1, 19.0%; pig 2, 16.9%; pig 3, 20.3%; pig 4, 20.0%; pig 5, 23.8%). Furthermore, the number of reads obtained from kappa and lambda-containing transcripts were similar (42.9% vs. 57.1%, respectively).

### IGK expression

A total of 159,721 reads corresponding to IGK and covering the entire variable region were obtained. 101,714 reads (64%) matched most closely to genes from the *IGKV1* family and 58,007 matched most closely to the *IGKV2* family (Fig. 1A). *IGKJ2* accounted for the vast majority of J gene usage (Fig. 1A). In contrast, *IGKJ3* and *IGKJ5* were not expressed and both *IGKJ1* and *IGKJ4* were expressed at very low levels (approximately 1% of all *IGKJ* genes). There was no discernable association between *IGKJ* and *IGKV* gene usage. Compared to the annotated germline sequences for *IGKV*, the median number of predicted amino acid mismatches varied from 3 for *IGKV2-6* to 10 for *IGKV1-11* (Fig. 1E). No major differences in *IGKV* gene usage were noted between individual pigs. Of the nine putatively functional *IGKV* genes, two (*IGKV1-7* and *IGKV1-14*) were not expressed in any of the five pigs.

**Figure 1.**
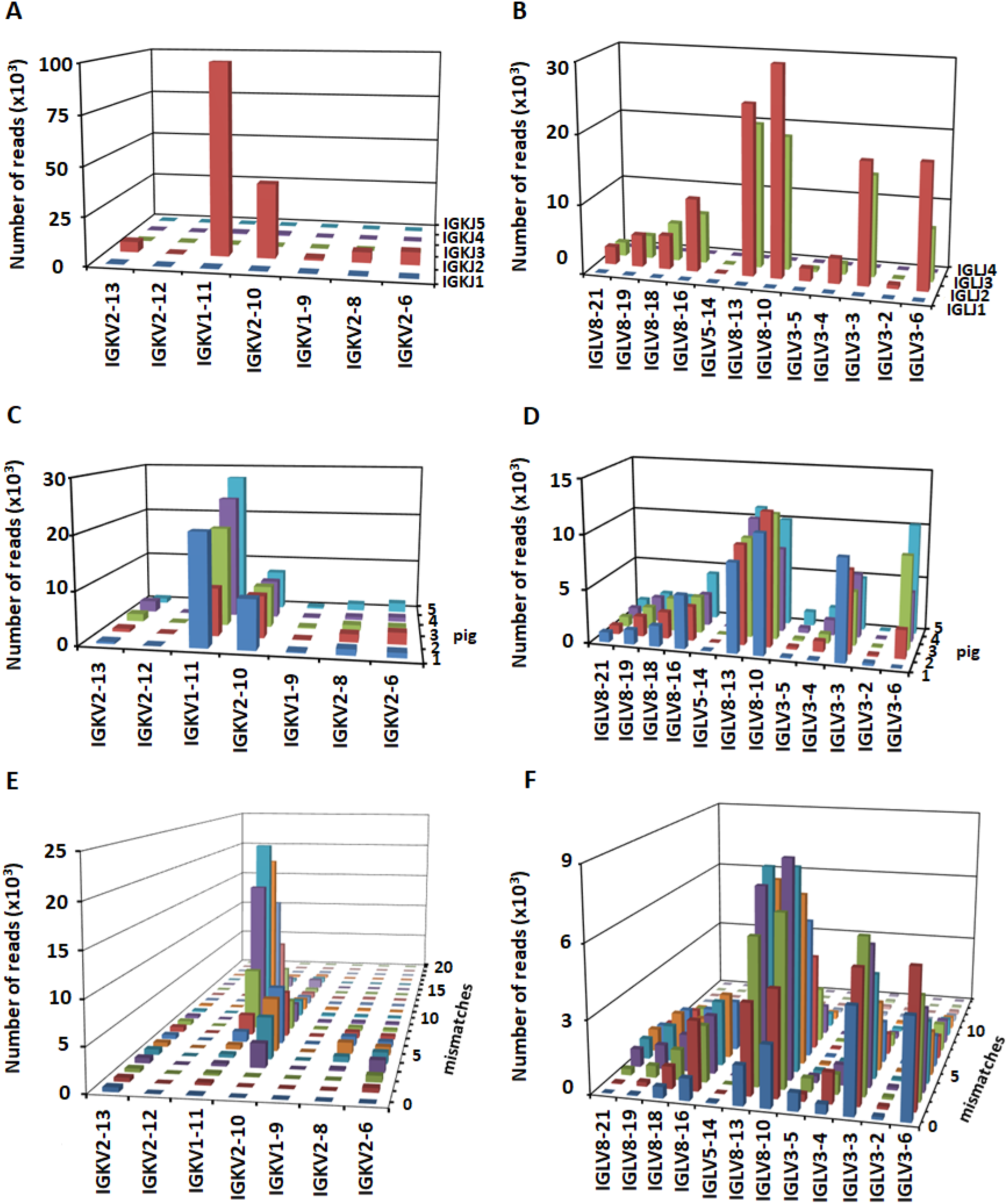
Light chain gene usage. *IGKV*-*IGKJ* (A) and *IGLV*-*IGLJ* (B) combinatorial usage. *IGKV* (C) and *IGLV* (D) gene usage in individual pigs. Deviation in amino acid sequence from the germline reference sequence for *IGKV* (E) and *IGLV* (F).

### IGL expression

A total of 212,419 full-length lambda variable region reads were obtained. *IGLV* gene BLAST results were dominated by *IGLV8* family members (67.6%), followed by *IGLV3* family members (32.4%), and low expression (0.03%) of *IGLV5-14* also was observed (Fig. 1B). *IGLJ* gene usage was divided between *IGLJ2* (59% of total reads) and *IGLJ3* (41% of total reads), and neither *IGLJ1* nor *IGLJ4* were expressed (Fig. 1B). There was no association between specific *IGLV* and *IGLJ* genes, as was observed for IGK. One apparently functional gene, *IGLV2-6*, showed no expression in any of the five animals, even though a conserved forward primer specific for the *IGLV2* leader region was used for PCR amplification (Table 1), and post-PCR purification of a gel slice in the expected size range was included in the amplicon pool that was sequenced. The PCR amplifications for the pseudogene *IGLV8-21* typically yielded a faint band of appropriate size. However, our sequence analysis failed to identify transcripts from this gene, suggesting that these products were non-specifically amplified transcripts from other members of the *IGLV8* family.

Of the *IGLV3* genes, *IGLV3-3* and *IGLV3-6* are the most highly expressed, yet this expression varies substantially between individuals. Pig 1 is known to be homozygous for a deletion of *IGLV3-6* and pig 2 likely possesses a different allele (12). In these animals, *IGLV3-6* transcripts were therefore either absent entirely or expressed at a relatively low level, respectively, compared to the other individuals. In contrast, *IGLV3-3* expression was higher in pigs 1 and 2 compared to the others (Fig. 1D). The usage of *IGLV3-6* was variable between the five pigs (12). Interestingly, in pig 1, which was previously found to completely lack this gene, *IGLV3-3* was almost exclusively the only *IGLV3* family member expressed (Fig. 1D). In the other animals found to possess a functional copy, however, *IGLV3-6* was one of the most highly expressed genes in the light chain repertoire (Fig. 1D).

### Light chain CDR3 length variation

Although a hallmark of heavy chains, CDR3 length variability is typically low in light chains, due to the lack of terminal deoxynucleotidyl transferase (TdT) activity during light chain rearrangement (22). However, we did observe some variation among the putatively functional light chain transcripts. Approximately 12.5% of IGK CDR3s differed in length from the germline encoded 9 amino acids, with a range of 4 to 16 amino acids. In contrast, 33.5% of IGL CDR3s differed from the canonical 10 or 11 amino acids, ranging from 4 to 23 amino acids in length (Fig. 2).

**Figure 2.**
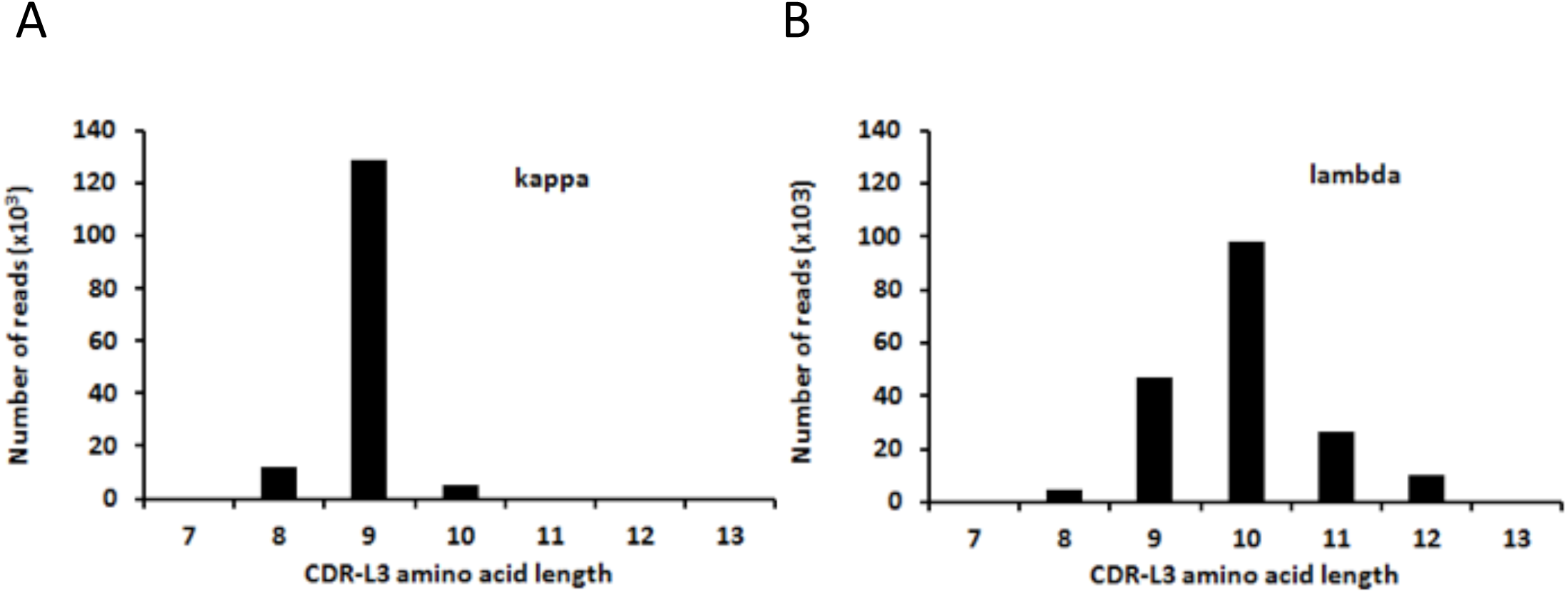
Length distributions of pig light chain CDR3. (**A**) Distribution of IGL CDR3 from 187,810 reads. (**B**) Distribution of IGK CDR3 from 147,189 reads.

### Diversity and richness of the IGK and IGL repertoires

Log-log abundance curves indicate a power-law distribution in which the vast majority of light chain sequences are exceptionally rare and a small number are very common (Fig. 3). The most abundant clones represented >1% of the total repertoire, while more than 50% of all sequences were singletons (IGK: 51.2±5.85%; IGL: 55.4±6.96% (mean±1 standard deviation)). Chao1 repertoire richness was estimated using deduced amino acid sequences for the full length V-J region from each animal. Lower bound richness estimates of 1.6 × 10^5^ to 2.8 × 10^5^ molecules per pig were obtained for the IGL repertoire and estimates of 1.1 × 10^5^ to 1.7 × 10^5^ molecules per pig were obtained for the IGK repertoire (Fig. 4). Their sum approximates the estimated richness of the porcine heavy chain VDJ (∼2.7×10^5^; Schwartz and Murtaugh, unpublished data).

**Figure 3.**
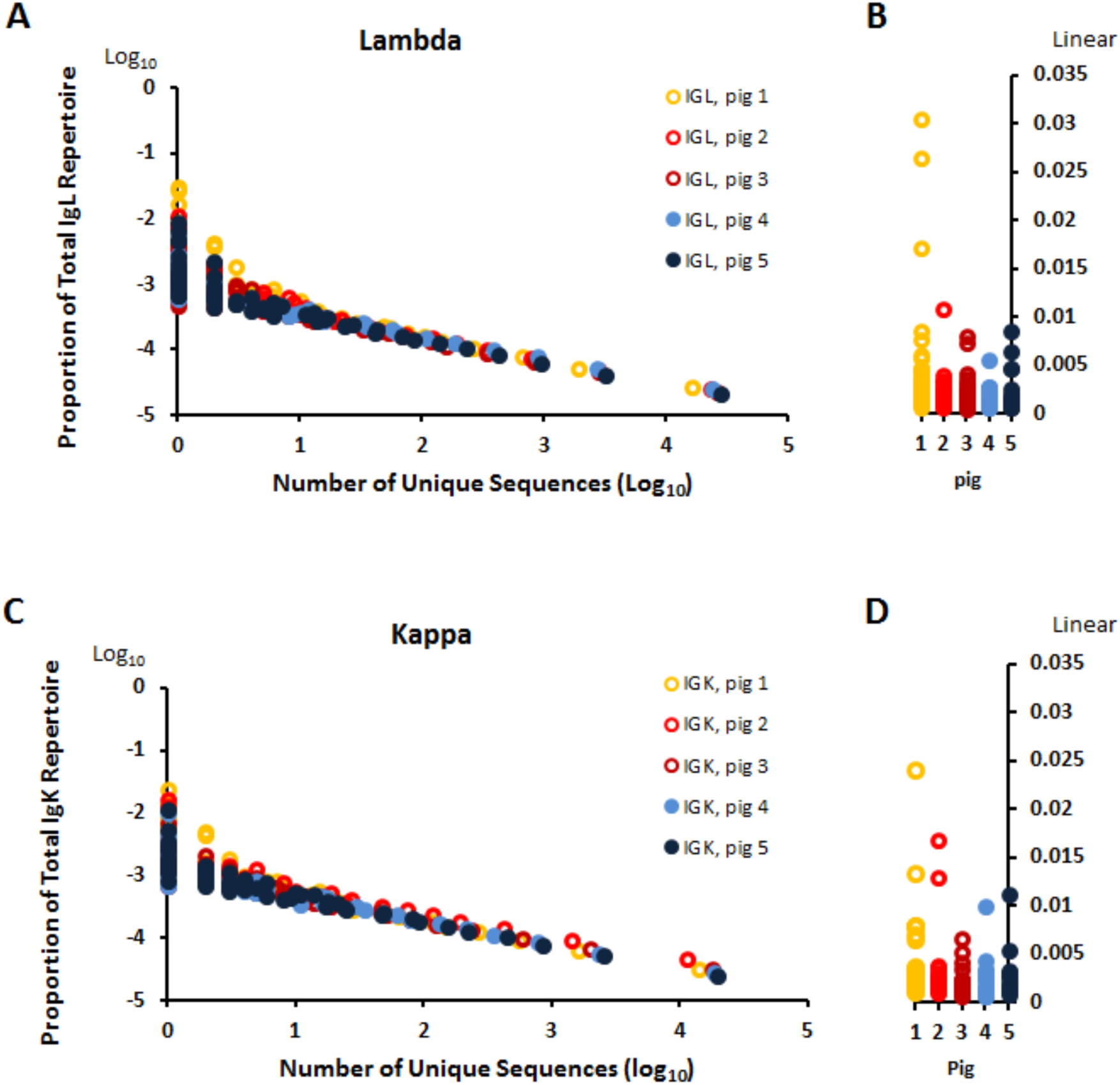
Light chain abundance distributions in PRRSV-infected (open circles) and uninfected (closed circles) pigs. (A) Relative abundance of lambda sequences as a proportion of the total lambda repertoire. (B) Effect of PRRSV infection on the abundance of unique lambda clones (data on Y-axis of panel A, shown on a linear scale). (C) Relative abundance of kappa sequences as a proportion of the total kappa repertoire. (D) Effect of PRRSV infection on the abundance of unique kappa clones (data on Y-axis of panel C, shown on a linear scale).

**Figure 4.**
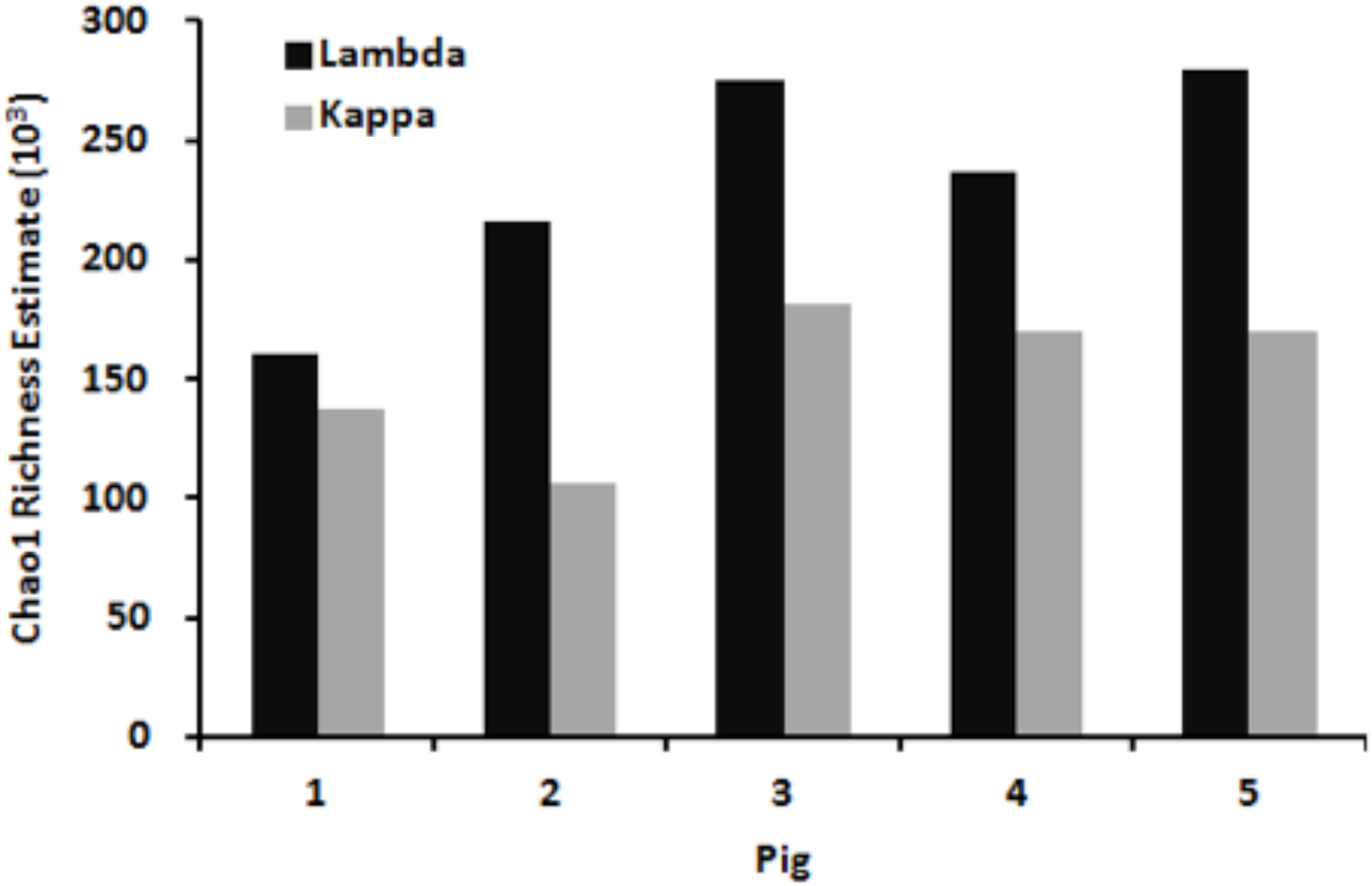
Lower-bound Chao1 richness estimates for the total size of the light chain repertoire among individual pigs. Lambda: average, 2.3×10^5^; standard deviation: ±4.9×10^4^. Kappa: average, 1.5×10^5^; standard deviation, ±4.9×10^4^.

The presence of dramatic allelic variation in some V genes, particularly for *IGLV3-6*, suggests that framework variation in the *IGLV* and *IGKV* family members might contribute to light chain repertoire diversity (12). Phylogenetic analysis showed high levels of nucleotide sequence variation within and between expressed light chain family members, varying by up to 11% and 15% pairwise differences within *IGKV1* and *IGKV2*, respectively, and 8% and 28% in *IGLV8* and *IGLV3*, respectively, while the highest pairwise similarity between family members varied from 43% (*IGKV1* versus *IGLV5*) and 62% (*IGLV3* versus *IGLV8*) (Fig. 8). The framework region of *IGLV3-3* is also distinct from the other *IGLV3* family members (Fig. 5). Within each group, variation is present in framework and CDR regions, with only the CDR3 region showing extensive variation, in part because of CDR3 length variability.

**Figure 5.**
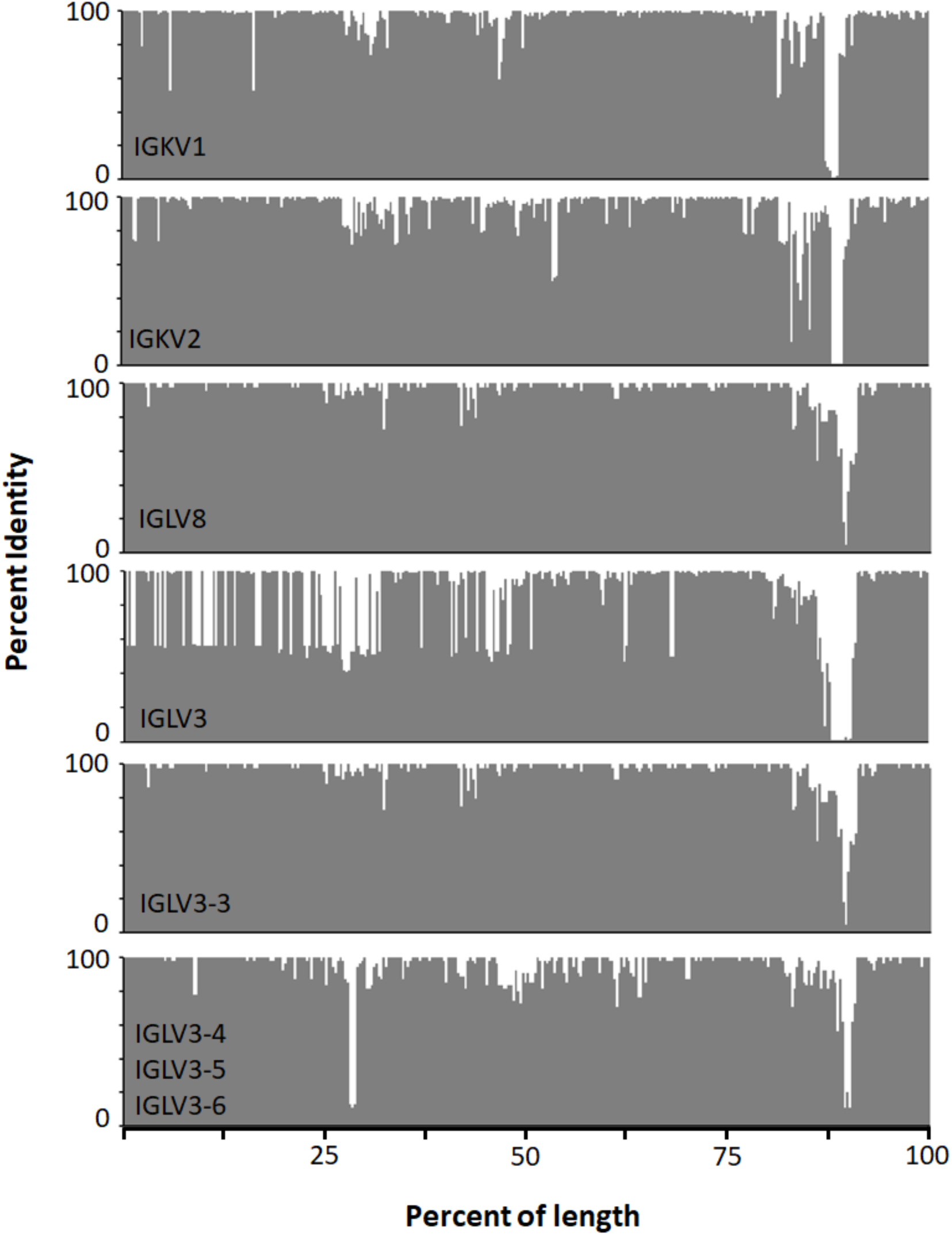
Distribution of amino acid sequence variation across light chain VJ region. The distribution for *IGLV3* includes *IGLV3-3, IGLV3-4, IGLV3-5*, and *IGLV3-6* for which the former is also shown separate due its greater genetic distance from the other three genes.

### The anti-PRRSV response

PRRSV infection caused a shift in light chain abundance distributions toward fewer, more abundant light chain sequences. Individual clonal populations were identified that accounted for one to three percent of the entire repertoires of individual pigs (Fig. 3). Individual variation in the abundance of specific clones is striking. The most abundant *IGLV*-*IGLJ* clone in this study, clone A in pig 1, was not present in uninfected pigs 3 and 4 and also was rare in PRRSV-infected pig 3 (Supplemental Table 1; Supplemental Table 2). The five most abundant clones in PRRSV-infected pig 1 were also present in infected pigs 2 and 3, but the five most abundant clones in PRRSV-infected pig 3 were absent or rare in the other two infected pigs. The five most abundant clones in pig 2 show somatic changes from the germline sequence, but 5 of the 10 most abundant clones in pigs 1 and 3 are unchanged from germline, the same ratio as in the dominant clones of the uninfected animals (Supplemental Table 1). Several highly abundant clones (e.g. clones B, E, L, M, and R) are present in multiple animals, including both PRRSV-infected and uninfected pigs (Supplemental Table 1). Amino acid sequence variation in abundant lambda clones was due primarily to mutations in both framework and CDR regions. The CDR1 region showed 4 sequence motifs with lengths of 5, 6 (n=2) and 9 amino acids, while CDR2 had no length variation and CDR3 length was 8±1 amino acids with one exception of 11 aa (Supplemental Table 2).

Unlike the IGL, abundant kappa clones in every case were mutated from the germline sequence, with an average of 7.4 amino acid differences in *IGKV* (Supplemental Table 3), compared to an average of 1.6 in lambda clones. These differences are nearly entirely mutational, as there were no length differences in CDR2 or CDR3, and only 3 length classes (6, 10, and 11 aa) in CDR1 (Supplemental Table 4). Kappa clones A and E were dominant in both PRRSV infected and uninfected pigs (Supplemental Tables 3). By contrast, pig 3 possessed clones I, J, and K which were not present in any other pig. Except for two *IGKV2-8* clones in pig 2, all abundant clones were derived from either *IGKV1-11* or *IGKV2-10* (Supplemental Table 3).

## Discussion

Unraveling of the mechanisms of antibody diversification in previous decades revealed combinatorial and mutational machinery able to produce a nearly limitless array of antigen-binding variability. Paradoxically, exposure of animal populations to an antigen consistently results in a wide range of response intensities. Therefore, the functional antibody repertoire of animals must be limited and variable among individual members of the population. Characterization of antibody repertoires and sources of individual variation in repertoire diversity in the absence and presence of antigenic challenge is essential to understand variation in antibody responses to infection and vaccination. Here we show that pigs, which are susceptible to a wide variety of microbial infectious diseases, use both IGL and IGK loci to produce repertoires in excess of 100,000 lambda and kappa light chains, even though V gene usage is limited and CDR complexity is low.

The IGK repertoire in the present study is dominated by the expression of only two genes, *IGKV2-10* and *IGKV1-11*, of which the latter is more highly expressed than the former. This result contrasts previous reports suggesting that *IGKV2* family members are more abundantly expressed, at least among preimmune piglets (8, 23). Two putatively functional V genes, *IGKV1-7* and *IGKV1-14*, lack a canonical octamer (ATCTGCAT) and both possess a non-canonical 5′ splice site in their intron (GT -> GG). As shown in Figure 2, there is a notably high level of sequence divergence in IGK transcripts from the germline annotation of the 14 most C-proximal *IGKV* genes in the locus (9). Since a high degree of heterozygosity was present, it is possible that individual diversity may be due to allelic variation, as has been shown in *IGLV* genes (11, 12). Allelic variation also has been shown in humans to significantly affect repertoire diversity (24, 25). However, whereas previous studies focused on heavy chain repertoire diversity and variation, we show here that light chain variation also makes a substantial contribution to overall antibody diversity using mechanisms that do not involve large CDR3 length variation and amino acid sequence replacement. Thus, generation of light chain repertoire diversity lies at the opposite end of the spectrum from bovine heavy chain diversity generation characterized by unusually long and complex CDR3 regions (26).

Diversity estimates as well as the raw sequencing results indicate that lambda expression accounts for 55 to 60% of total light chains, and consists overwhelmingly of *IGLV8* family members, followed by *IGLV3* family members. These results are consistent with previous reports of the pig lambda repertoire (10, 27), which also suggested that there are age-dependent changes in *IGLV* usage. Although putatively functional, *IGLV2-6* was not expressed in any of the five animals studied. Our previous analysis indicated that this gene lacks a conserved octamer in the promoter region (ATTTGCAT -> ATTTGTAT) and a conserved heptamer in the recombination signal sequence (CACAGTG -> TACAGTG), which together appear to account for its lack of functional expression (9). Both *IGKV* and *IGLV* showed limited CDR diversity but substantial framework variation that is most striking in the *IGKV3* family. The broad distribution of amino acid variation, encoded in the DNA, across the majority of the light chain VJ regions, creates extensive diversity and, presumably, in the structural stability of the antibody molecule. It also increases the likelihood that allelic variation will contribute substantially to variation in the antibody responses of individual animals to antigenic challenge.

Viral infection skewed light chain responses to more highly abundant clones which is suggestive of antigen-driven clonal expansion. It was previously argued that PRRSV induces a pronounced nonspecific B-cell activation and hypergammaglobulinemia that also could skew antibody responses to abundant, antigen-independent clones due to its persistence in the host. However, it is apparent now that PRRSV infections are resolved completely, and that evidence is lacking for hypergammaglobulinemia in conventional swine infected or vaccinated with PRRSV and reared under a variety of conditions (28). The specificity of highly expressed light and heavy chains is resolvable due to the ability to clone and express recombinant antibodies. Therefore, we expect that the antigen specificity of highly abundant clones will be determined in later studies.

Interestingly, pig 1 showed a marked reduction in total estimated richness in the lambda repertoire compared to the other four individuals. The reduction appears to be due to genetic constraints in the C-proximal *IGLV3* gene cluster. In particular, the sequences obtained from this animal indicated a high degree of IGL homozygosity. Within the C-proximal *IGLV* cluster, only *IGLV3-3* was expressed at high levels, while the other *IGLV3* genes were nearly unused. In addition, *IGLV3-6* expression was absent. As the most abundantly expressed antibody sequence from this animal was of *IGLV3-3* origin, it is possible that the reduction in richness was due to genetic constraints and not PRRSV infection. Taken together, the findings show that the light chain repertoire is equivalent in size and complexity to the heavy chain repertoire, in the range of 10^5^ to 10^6^ unique light chains per individual, with fully half of the sequences present as singletons. These richness estimates closely resemble the reported richness of the human light (1.6 × 10^5^) and heavy (2.2 × 10^5^) chain repertoires which were estimated using a capture-recapture rarefaction method (38).

Analyses of the human light chain repertoire has revealed significant differences between individuals in their CDR3 sequences and V gene usage (29, 30). Indeed, in a study consisting of four humans, approximately 60 percent of light chain rearrangements were present in more than one individual (30). Furthermore, approximately 20 percent of light chain CDR3s were shared by more than one individual in a study consisting of five humans (29). Our data also show substantial individual variation in the molecular distribution of light chains in pigs; the most dominantly expressed sequences in some animals are not present in others. These features are also characteristic of antibody heavy chains in other species such as zebrafish (31, 32). However, since zebrafish have only ∼300,000 B-cells and a heavy chain richness of 1200-3700 unique antibodies per individual (31, 32), the actual diversity of antibodies produced in a population will be distributed stochastically among individuals such that broad variation in ability to respond to antigens will be a feature of the population more than the individual. The spleen of swine and humans, alone, has a mass more than 100 times larger than a zebrafish (∼150 g versus <1 g, (33-36)), and with many additional lymphoid tissues, the total B-cell compartment of humans and large animals is orders of magnitude larger than a zebrafish or mouse, whose spleen weighs about 100 mg (37). Therefore, swine are a relevant model of human antibody diversification in which the immune response capacity of individuals is critical to understanding variation in the effectiveness of immunity to pathogens.

## Acknowledgements

We thank Juan Abrahante for assistance and advice in sequence data management and Gayathri Dileepan for technical assistance. Funding was provided by the National Pork Board grant 10-139 (J.C.S. and M.P.M.). J.C.S. was supported by the Molecular Virology Training Grant T32 AI83196 from the National Institutes of Health, a Doctoral Dissertation Fellowship from the University of Minnesota, and is currently supported by the United Kingdom Biotechnology and Biological Sciences Research Council (BBSRC) project BBS/E/I/00001710, “Exploring the impact of genetic variation on the livestock immune response to pathogens.”

